# Drivers of temporal bias in biodiversity recording by citizen scientists

**DOI:** 10.1101/2024.07.22.604598

**Authors:** Inês.T. Rosário, Patrícia. Tiago, Sergio Chozas, César Capinha

**Author notes:** Corresponding Authors: César Capinha | Inês T. do Rosário.

## Abstract

1. Citizen science data is increasingly important for ecological research, biodiversity conservation and monitoring. However, these data often suffer from biases due to uneven recording efforts by citizen scientists. Biases caused by intra-annual differences in recording activity can be particularly severe, hindering the use of citizen science data in research areas such as population dynamics and phenology. Therefore, understanding the factors driving recording activity is essential.
2. In this study, we provide a detailed assessment of how weather and calendar-related factors influence biodiversity recording activity by citizen scientists at a daily resolution. To perform this, we analyse the recording patterns for six tree species in the Iberian Peninsula, which maintain a fairly consistent appearance throughout the year. Observation data were collected from iNaturalist, a leading platform for citizen-science data. We used boosted regression trees (BRT) to compare observed recording activity patterns with those expected by chance. Our analysis included a comprehensive set of explanatory variables, such as day of the week, month, holidays, temperature, accumulated precipitation, wind intensity, and snow depth.
3. The BRT models effectively identified the drivers of recording activity, with the correlation between predicted and observed temporal patterns (left out of model training) ranging from 0.47 to 0.96, depending on the species. The day of the week, daily temperature, and month of the year consistently emerged as the main drivers. Recording activity was higher on weekends, to some extent on Fridays, and during the spring months. Extreme low and high temperatures generally correlated with lower recording activity, although there were exceptions. Wind speed and precipitation had a moderate influence, with higher wind intensity and accumulated precipitation leading to decreased activity. Holidays and accumulated snow had very minor relevance across species.
4. Our findings show that citizen scientists record more frequently on weekends, during mild weather, and in spring. By addressing these biases, we can maximize the utility of citizen-collected data for research and applied purposes, ensuring robust and reliable conclusions that enhance ecological understanding and conservation efforts.

## 1. Introduction

Currently, citizen science is an invaluable source of biodiversity data (Callaghan et al., 2021). These data, recorded mostly by non-experts and often empowered by specialised smartphone apps, are now among the most abundant available for several taxonomic groups (Groom et al., 2017; Kelling et al., 2019). Thus, citizen science data are crucial for supporting assessments of the current state of biodiversity (Chandler et al., 2017), monitoring the spread of invasive species (Gallo & Waitt, 2011; Johnson et al., 2020), and tracking the distribution and population trends of species (Chandler et al., 2017; Dennis et al., 2017; Horns et al., 2018). Additionally, citizen science initiatives play a significant role in ecological research, conservation planning, and policymaking by providing large-scale, geographically diverse datasets that are otherwise difficult and expensive to obtain (Hochachka et al., 2012; Callaghan et al., 2019). Furthermore, the engagement of the public in scientific research fosters a greater awareness and understanding of environmental issues, promoting conservation efforts at a community level (Bonney et al., 2016, Chozas et al., 2023).

While citizen science has significantly advanced biodiversity data collection and environmental research, it also presents several important biases and analytical challenges. These include spatial biases that influence the geographic distribution of observations, taxonomic biases that result in uneven representation of different species or groups and temporal biases that affect the timing and seasonality of data collection (Bird et al., 2014; Tiago et al., 2017; Troudet et al., 2017). Temporal biases are particularly relevant, owing to strong variation in levels of recording activity by citizen scientists (Courter et al., 2013; Bird et al., 2014; Backstrom et al., 2024). This variability results from factors that influence the willingness and availability of citizens to engage in recording activities (Hobbs &White, 2012; Peter et al., 2021), ultimately hindering the effective use of collected data to analyse the temporal dynamics of the recorded phenomena. Although several analytical strategies have been proposed to address this issue (Bird et al., 2014, Dennis et al., 2013; Gonsamo & D’Odorico, 2014; Kosmala et al, 2016; Weiser at al., 2020), challenges persist, particularly in researching seasonal phenomena that depend on high temporal resolution data, such as population dynamics and phenology (Callaghan et al., 2019; Primack et al., 2023). Therefore, thoroughly evaluating these biases and understanding their underlying causes is crucial for improving the comprehensive and accurate use of citizen science data. By accounting for these biases, the reliability, and validity of conclusions drawn for temporally-detailed assessments of ecological phenomena can be significantly enhanced (Mora et al., 2024; Tiago et al., 2024b).

Variations in levels of biodiversity recording activity by citizen scientists throughout the year reflect the interplay of two components. The first component pertains to the seasonal dynamics of the actual ecological phenomena being recorded. Variations in the timing and magnitude of these phenomena, such as plant blooming, butterfly emergence in the imago stage, or bird and insect migration events, inherently determine the opportunities for citizens to record them (e.g., Monarch Larva Monitoring Project, Brenskelle et al., 2021). The second component involves factors external to the ecological phenomena determining citizen scientists’ availability and willingness to participate, such as the day of the week, public holidays, or the time of year (Courter et al., 2013; Di Cecco et al., 2021), as well as weather conditions. A common aim of researchers using citizen science data to study ecological dynamics is to ’remove’ the influence of the later component on the temporal patterns of recording activity (e.g., Capinha et al., 2024). By doing so, the resulting (‘unbiased’) patterns more accurately represent the true dynamics of ecological phenomena. However, this is often challenging because the effects of both components are intertwined, hindering the interpretation of observed trends in the data.

Previous studies have assessed the role of external factors in driving levels of activity of biodiversity recording by citizen scientists. The factors assessed are predominantly calendar-related, such as the identified weekend bias and the week-of-the-year bias (Courter et al., 2013; Di Cecco et al., 2021; Díaz-Calafat et al., 2024). Additionally, the impact of citizen-science-specific events, such as ’BioBlitzes’ or the ’City Nature Challenge’, has also been investigated (Di Cecco et al., 2021; Tiago et al., 2024). However, most of these assessments were based on the raw number of records contributed by citizen scientists (e.g., Di Cecco et al., 2021; Díaz-Calafat et al., 2024), overlooking how variations in the timing and magnitude of the ecological phenomena themselves influence recording patterns. Moreover, to date, there has been no comprehensive assessment of the role of weather-related factors, despite the well-known influence of weather on the willingness to engage in outdoor activities (Tucker & Gilliland, 2007). Weather-related factors are plausibly relevant drivers of biodiversity recording, potentially more so than calendar-based ones, and their effects may be complex. For instance, some meteorological variables might be expected to have a monotonic effect, such as a consistent decrease in outdoor activity with increasing precipitation. On the other hand, peaks of recording activity plausibly occur at moderate rather than extreme low or high temperatures. Ultimately, it is essential to understand the joint effects of calendar and weather-related factors on the levels of activity of citizen scientists. This understanding must be developed using approaches that are robust to the seasonal variations of ecological phenomena.

Here, we provide a detailed assessment of how calendar-related and weather factors influence the levels of biodiversity recording activity by citizen scientists. We focus on the recording patterns of a set of evergreen tree species that fulfil several key criteria: conspicuousness, ease of identification, and, most importantly, consistent appearance throughout the year. Our core assumption is that due to minimal changes in their appearance, the temporal patterns of recording for these ’benchmark’ taxa primarily reflect variations caused by external factors. We analyse observation records from the Iberian Peninsula and use a comprehensive set of explanatory variables representing both calendar and weather-based factors, including the day of the week, month, bank holidays, temperature, accumulated precipitation, wind intensity, and snow depth. By evaluating the relative importance of each variable and their associations with levels of recording activity, we offer a thorough assessment of the external factors influencing biodiversity recording by citizen scientists.

## 2. Methods

### 2.1 Species occurrence data

The analysed species observation data were collected from the iNaturalist platform (www.inaturalist.org). iNaturalist is a nonprofit social network designed for naturalists, citizen scientists, and biologists, centred around the concept of mapping and sharing biodiversity observations globally. iNaturalist is accessible via its website and mobile applications. It features an automated species identification tool and fosters a collaborative approach where users collaborate in identifying organisms from photographs or sounds (www.inaturalist.org/pages/about).

From iNaturalist we collected biodiversity observations from seven evergreen tree species. These were five pines: *Pinus halepensis* (Aleppo pine), *P. nigra* (Austrian pine), *P. pinaster* (maritime pine), *P. pinea* (stone pine) and *P. sylvestris* (Scots pine), and two oaks: *Quercus rotundifolia* and *Q. suber* (holm and cork oak, respectively). We selected these species because they are the pine species and evergreen oaks that are more widely distributed in the Iberian Peninsula and exhibit minimal variation in their appearance through the year. Our core assumption is that, due to reduced seasonal changes in their appearance through the year, the temporal patterns in recording of these taxa should largely reflect variations in recording activity itself (cf. Capinha et al., 2024). This should be particularly the case with pines, which maintain a similar appearance throughout the year. They grow new leaves as they shed old ones, and their (female) pine cones take longer than a year to mature (e.g., one and a half years for *P. nigra,* two years for *P. halepensis and P. pinaster,* and three years for *P. pinea*), resulting in cones of various ages and sizes throughout the year (Earle, 2018). For the two oak species, while they are also evergreen and maintain a constant appearance of their foliage and branches, during Autumn they produce acorns (Bonal & Muñoz, 2007, Pons & Pausas, 2012). The acorns may imply, to some extent, a slight increase in attractiveness for recording during this period. Interpretation of results for these taxa (below) take this possibility into account.

The observation data collected from iNaturalist file included: geographical coordinates of location of observation, date of observation, username of observer, species’ scientific name, and quality grade. We filtered these data and kept only the records supplying geographical coordinates, full date (year, month, and day) and a research quality grade (i.e., species identification confirmed by at least two identifiers). To avoid years in which observation recording may have been affected by anomalous factors, the two years of lockdown due to the Covid-19 pandemic were excluded. In addition, the number of records made prior to 2017 was very reduced. Therefore, the observation data collected represented the periods from 2017 to 2019 and 2022 to 2023.

### 2.2 Assembly of dependent and predictor data

To compile the data for each species for analysis, we began by retaining only a single record per combination of date and 0.1° ⨉ 0.1° (approx. 10 x 10 km) grid cell, corresponding to the spatial resolution of the predictor data (see below). This step aimed to avoid including multiple records from the same recording event, such as a single recorder submitting multiple records of the same specimen (i.e., ’duplicates’). A total of 3,532 observation records were kept for *Q. suber*, 4,744 for *Q. rotundifolia*, 938 for *Pinus sylvestris*, 1272 for *P. halepensis*, 2,118 for *P. pinea* and 1,246 for *P. pinaster*. *P. nigra* was excluded from the analysis because there were few observations (128).

Next, for each species, we generated an equal number of records with the same geographical coordinates and year of observation but with randomly generated day and month values. These records are intended to represent the temporal distribution expected if recording events were randomly distributed throughout the year, referred to as ’temporal pseudo-absences’ (Capinha et al., 2024). We then combined the observed records (coded as ’1’) with the temporal pseudo-absences (coded as ’0’) into a single data set. Thus, for each species, we created a data set containing both the actual observation data and data representing the temporal distribution of records if observations were made on random days.

Each record in these data sets was then characterised by a comprehensive set of calendar and weather-related variables, which are expected to be the main drivers of individuals’ willingness to go outside and observe and record biodiversity. The calendar variables included the day of the week, the month, and whether the day was a bank holiday. The first two variables were calculated directly in R (R Core Team, 2024) using base functions, while holidays were identified manually from the website Timeanddate (www.timeanddate.com). These calendar variables were selected based on earlier literature assessing the influence of the day of the week and the time of year on species recording (Courter et al., 2013; Di Cecco et al., 2021). Additionally, we considered that bank holidays could also be a relevant factor in determining the propensity for biodiversity recording.

The weather variables included the mean temperature of the day (°C*10), total accumulated precipitation of the day (mm), mean wind speed on the day (m/s), and snow cover depth on the day (cm). These values were sourced from ERA5Ag at 0.1° ⨉ 0.1° resolution (Boogaard at al. 2020). We selected these weather variables based on previous studies assessing factors that influence outdoor activities, with temperature and precipitation being the most commonly considered (Verbos et al., 2018). Wind and snow may also be relevant factors, impacting, for example, people’s outings to natural parks, particularly in higher-altitude areas (Verbos and Brownlee, 2017).

### 2.3 Data analysis

To identify predictors of temporal variation in recording effort, we used boosted regression trees (BRT) (Elith et al., 2008; Hijmans et al., 2017). BRTs are ensembles of decision trees, where each tree is fitted iteratively to reduce the prediction errors of the ensemble. This algorithm effectively combines multiple weak learners (decision trees) to create a robust predictive model (Elith et al., 2008). We implemented BRT models in R using the ‘gbm.step’ function from the ’dismo’ package (Hijmans et al., 2017). This function supports automated tuning of the optimal number of trees to include in the ensemble through internal cross-validation. Additional important hyperparameters to consider for tuning are tree complexity (tc) and learning rate (lr). Tree complexity indicates the maximum number of interactions in each tree, while the learning rate determines the contribution of each tree to the overall model. Hence, for each species, we tested combinations of common values of learning rates (0.0001, 0.0005, and 0.001) and tree complexities (3 and 6). To evaluate model performance with each combination, we randomly set aside 30% of the data for comparison with predictions. Level of agreement between left-out data and predictions was measured using the Boyce index (Hirzel et al., 2006). Applied to our models, this index measures how well the predictions of species recording correlate with the temporal distribution of the actual observation records. We perform these calculations with the ‘ecospat.boyce’ function from the ’ecospat’ package (Di Cola et al., 2017). The values corresponded to Pearson correlation coefficients, ranging from -1 to 1, with higher positive values indicating better model performance. For our data, the Boyce index is preferable as a performance metric compared to discrimination-based metrics such as the area under the receiver operating characteristic curve (AUC). This is because discrimination-based metrics evaluate the model’s ability to correctly predict both classes in the dependent variable. However, in our case, pseudo-absences are randomly generated over time, including periods favourable for biodiversity recording. Therefore, considering the model’s ability to predict these records would incorrectly deflate their performance.

After identifying the best-performing set of parameters for each species, we then fitted a model using the full data set. From each of the full models, we obtained the relative importance of each variable. These importances are based on the number of times a variable is selected for splitting, weighted by the improvement to the model resulting from each split (Friedman & Meulman, 2003). This is a robust and widely used method in ecology to assess the influence of predictor variables (e.g., Elith et al., 2008). Additionally, we assessed how variation in the values of each predictor relates to the propensity for biodiversity recording. This was performed by extracting partial dependence plots, which show the effect of a variable on the response after accounting for the average effects of all other variables in the model.

## 3. Results

### 3.1 Model performance

The predictive power of the models varied with the combination of different values of learning rate and tree complexity (Table 1). However, the best-performing models achieved strong to very strong Pearson correlation values between the predictions of species recording effort and the periods when the species were effectively recorded (min r = 0.47 and max r = 0.96).

**Table 1.** Results of Boyce index for boosted regression models using distinct combinations of learning rate (Lrate) and tree complexity (TC). Boyce index values range from -1 to 1, with higher positive values indicating better model performance. Best performing models for each combination are shown in bold.

| Lrate | 0.001 | 0.001 | 0.0005 | 0.0005 | 0.0001 | 0.0001 |
| --- | --- | --- | --- | --- | --- | --- |
| TC | 3 | 6 | 3 | 6 | 3 | 6 |
| *Pinus halepensis* | 0.122 | **0.472** | 0.116 | 0.443 | 0.018 | 0.177 |
| *Pinus pinaster* | 0.66 | -0.592 | **0.743** | 0.471 | -0.494 | 0.536 |
| *Pinus pinea* | 0.528 | **0.812** | 0.25 | 0.611 | 0.613 | -0.04 |
| *Pinus sylvestris* | **0.757** | 0.67 | 0.684 | 0.722 | 0.591 | 0.574 |
| *Quercus rotundifolia* | 0.585 | **0.914** | 0.758 | 0.832 | 0.514 | 0.758 |
| *Quercus suber* | 0.836 | 0.752 | 0.713 | **0.951** | 0.425 | 0.848 |

### 3.2. Predictors of recording effort

Across species, three variables consistently emerged as the most important predictors of timing of recording: ‘weekday’ (median relative importance = 25.1%), ‘temperature of the day’ (24.6%) and ‘month’ (22.4%). Notably, depending on the species, there is also some interchange in the ranking of these variables. Following these three main predictors are ‘wind speed’ and ‘precipitation’, with lower but still relevant contributions (median importance = 14.3% and 8.4%, respectively). On the other hand, the ‘holiday’ predictor achieved negligible to low importance across species (median value = 0%; maximum = 4.5%), while ‘snow depth’ showed consistently negligible importance (median value = 0%; maximum = 0.01%).

Partial dependence plots describe the form of relationship between the probability of species recording and predictor values (Fig. 2). For ‘weekdays’, there is a consistent pattern of higher probabilities of species recording on Saturdays and Sundays compared to weekdays. Additionally, there is also a noticeable higher propensity for species recording on Fridays compared to remaining non-weekend days.

**Fig. 1 -.**
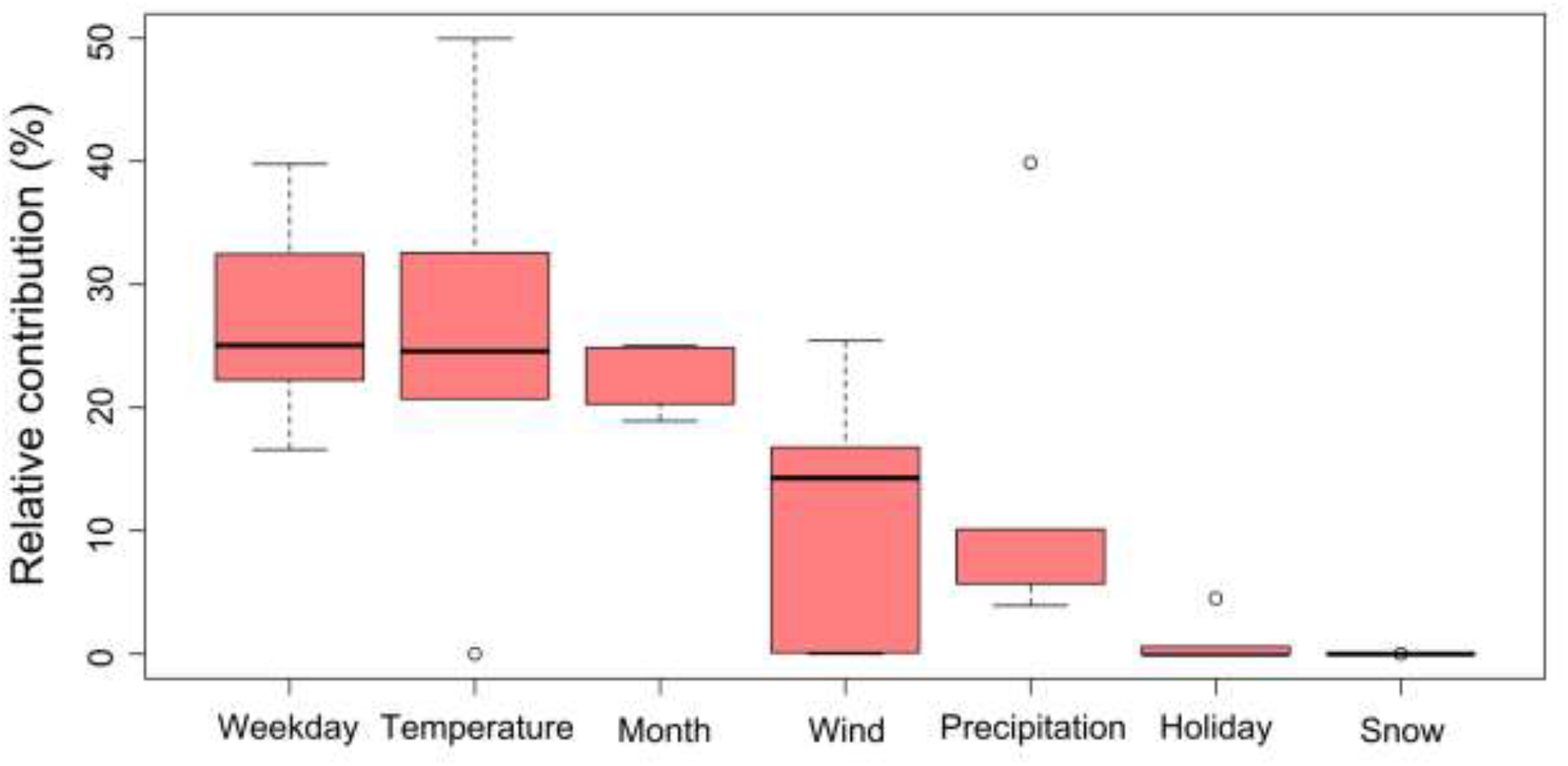
Relative importance of predictor variables used for predicting the timing of recording of benchmark taxa. Boxplots represent the variation of relative importances across the six species analysed. Higher values indicate a higher relative importance of the variable in distinguishing between the timing of records of the benchmark taxa and the timing of records expected randomly over time.

**Fig. 2 -.**
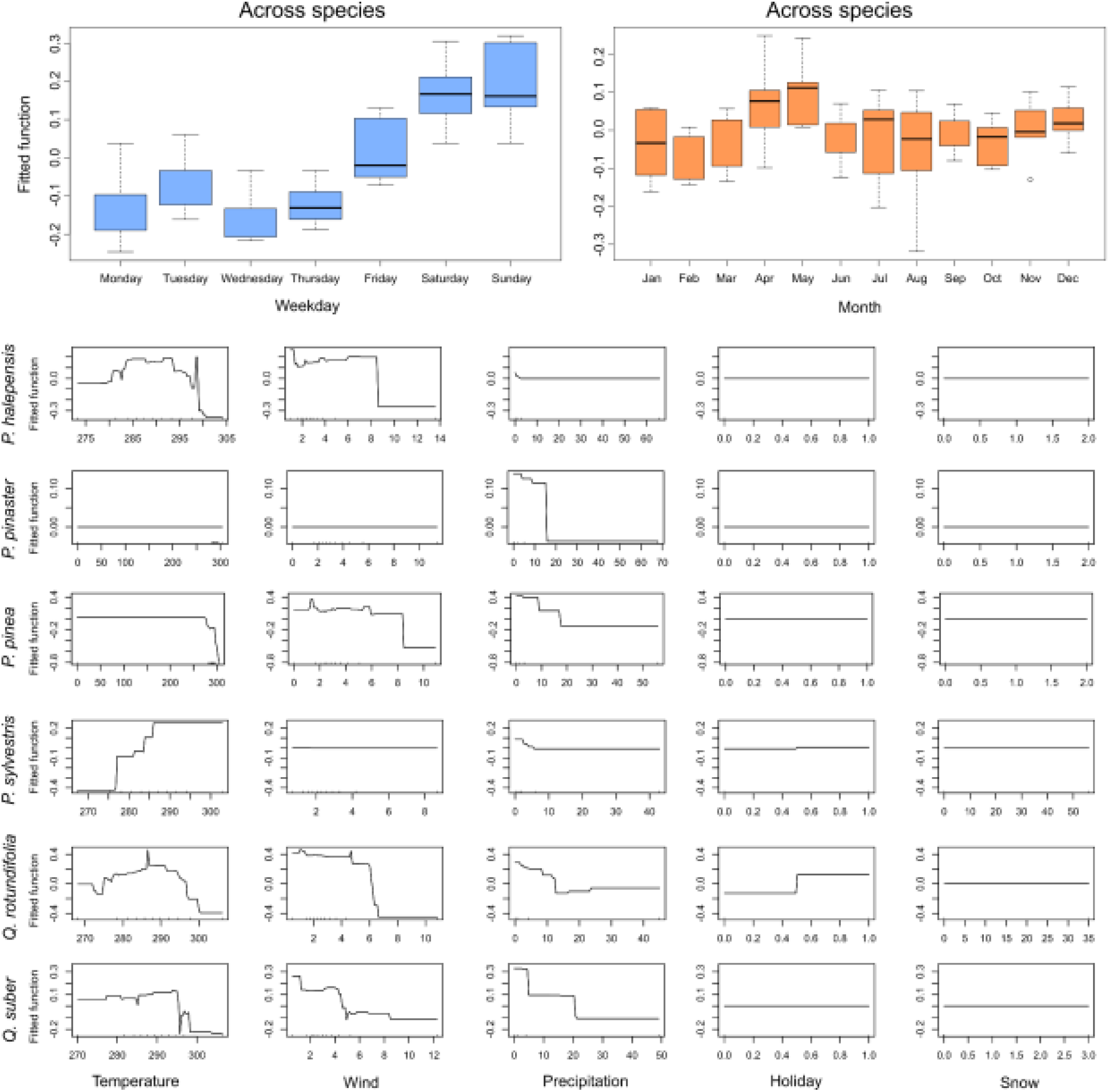
Partial dependence plots of variables used for predicting the timing of species observation recording. The top two panels represent the predictors ‘weekdays’ and ‘months’, sharing plot axes across models and depicting the variation of fitted values across the six benchmark species. The lower panel displays the remaining variables, each with distinct values in the *x* axis, and shows the fitted functions individually for each model.

For the ‘months’ predictor, the responses are more varied, with only two months, April and May, consistently showing higher propensities for species recording. For the two *Quercus* species there was no evidence of higher recording activity in the autumn months, when acorns are produced (Fig S3; supplement). Concerning ‘temperature’, distinct general trends can be observed.

For most species (excluding *P. pinaster* and *P. pinea*), models indicate a slight to sharp increase in recording activity with decreasing temperatures. However, most species also show a sharp decrease at extreme high temperatures. In contrast, *P. sylvestris* exhibits a consistent positive relationship between temperature and recording activity across the entire temperature gradient.

For ‘wind speed’, there is a largely consistent negative relationship, with increasing wind speed values associated with lower recording probability. The same general negative relationship is consistently found for precipitation. Holidays showed no apparent relationship for most species, reflecting their negligible importance. However, an expected positive relationship is still noticeable for two species (*P. sylvestris* and *Q. rotundifolia*). ‘Snow depth’ showed no apparent relationships, in line with the negligible relative importance of this predictor across species.

## 4. Discussion

In this study, we analysed the temporal patterns of citizen science recordings for a set of species with minimal phenological changes throughout the year. Using nearly 14,000 citizen-science observations of these benchmark taxa across the Iberian Peninsula, we found that the day of the week, mean temperature of the day and month of the year have a major influence in the levels of recording activity. Mean wind intensity and accumulated precipitation also showed relevant, albeit slightly lower, influences. These findings offer novel and deeper insights into the temporal factors driving citizen science recording activity.

Weekday was identified as the most influential variable across species, with Saturdays and Sundays (and to a lesser extent, Fridays) having higher levels of recording activity. This "weekend effect" has been previously identified, particularly in studies focused on birds (Fraser, 1997; Surmacki, 2005; Sparks, 2008). Our findings can be easily explained by the greater availability of people to engage in citizen science activities, as most do not work on weekends (Perry-Jenkins & Gerstel, 2020). Weekends serve a crucial social function by providing shared leisure time for families or groups of adults (Bryce, 2020). Activities such as BioBlitzes exemplify how weekends facilitate collective engagement in outdoor and educational activities while promoting a higher number of records on these days (Di Cecco et al., 2021). These events are typically organised to record as many species as possible within a specific period and are usually held on weekends to ensure greater community participation (e.g.: Meeus et al. 2023, Tiago et al. 2024b).

The time of year also had a relevant influence in levels of recording activity, with a notable increase during April and May. This pattern aligns with findings by Di Cecco (2021), who observed similar trends in global iNaturalist record submission data. These months correspond to spring in the study area, when the end of the colder season likely encourages more outdoor engagement. These months correspond to spring in the study area, marking the end of the colder season, likely encouraging people to engage in outdoor activities (Tucker & Gilliland, 2007). Spring is also particularly attractive for observing certain taxonomic groups or particular phenological events, such as the arrival of migratory birds (Greenwood 2007), plants in bloom or the emergence of insects (Daru et al., 2018). Although the species in our data are not subject to these fluctuations, it is likely that observers going out to observe more attractive species during these months will also record other species. Organised special events, often held in the spring, such as Fascination of Plants Day (https://plantday18may.org/), the Spanish Flora Biomarathon, the Portuguese Flora Bioblitz, Invasive species week, BioBlitzes, may also influence recording activity (Márquez-Corro, 2021; Chozas et al., 2023; Tiago et al., 2024a).

Temperature was also identified as a main driver of recording activity levels, with cross-species relevance similar to the variables “weekday” and “time of the year”. We found that days with extremely high or low temperatures had a lower propensity for recording most of our benchmark species. These results are consistent with previous studies indicating that extreme temperatures affect human outdoor activity patterns (Chen & Ng, 2012). During very hot or very cold periods, citizens are less likely to engage in outdoor activities, such as wildlife observation and recording. One exception was found for *P. sylvestris*, where a monotonic positive relationship was found between daily temperatures and recording levels. This likely results from this species being mainly found in mountainous areas of the Iberian Peninsula (Earl, 2018), places where warmer temperatures should favour visitation.

Daily wind intensity and accumulated precipitation were also identified as important drivers of recording activity, though their effects were less pronounced than those of temperature. Increases in the values of both variables were associated with a reduction in species records. These results align with expectations based on studies of people’s predisposition for outdoor activities. Precipitation is widely regarded as the most adverse condition for outdoor activities (Steiger et al., 2016; Wagner et al., 2019), and although wind is less frequently mentioned, it can still impact people’s perception of temperature and increase discomfort, particularly at lower temperatures (Andrade et al., 2010). There are particular cases where extreme weather conditions can be attractive to naturalists, such as experienced birdwatchers seeking rare species brought by storms (Tryjanowski et al., 2023). However, this behaviour does not apply to the majority of cases, especially for the benchmark taxa we assessed, which are specifically used to evaluate citizen scientists’ responses to factors external to the recorded phenomena.

The other meteorological variable considered, snow depth, had no identifiable influence on the recording patterns of the species. This is likely because snow cover in the Iberian Peninsula is relatively limited in both spatial extent and temporal duration, thus reducing its potential impact on recording activities. In other locations, such as Northern Europe, where snow covers large regions for extended periods of the year, this factor could have a relevant and more pronounced effect. However, snow depth has been previously identified as having no significant influence on outdoor activities except for very specific cases that require snow, such as skiing or ice skating (Spinney and Millward, 2010).

It is somewhat surprising that no holiday effect was found in the recording of most species studied, considering that the reasons justifying higher recordings on weekends could also apply to holidays. Similarly, Knape et al. (2022) observed a weekend effect but found that holidays influenced only bird recordings, with no such effect for insects, fungi, or plants. These authors examined five taxonomic groups using data from the Swedish Species Observation System (Artportalen; https://www.artportalen.se/) to understand the effects of temporal patterns such as seasons, weekdays, and holidays. They found that only bird records increased during holidays, attributing this difference to a larger sample size for bird data or different observer communities. For instance, if most bird observers are retirees, the weekday effect diminishes, whereas if they are professionals recording as part of their work during the week, the effect is more pronounced. In our case, it is unlikely that the observers belong to different communities since all species are plants, specifically from the genera *Quercus* and *Pinus*. Another possibility is that holidays are often spent with family or friends and involve specific rituals, such as Christmas, Easter, Saints’ festivals, or Carnival (Santos et al. 2023; Vihalemm & Harro-Loit 2019), leaving little time for activities like biodiversity recording. Díaz-Calafat et al. (2024) found a negative relationship between the number of observations and holidays, particularly during the winter. These authors suggested the possibility of a confounding effect between the holiday and insect activity. This does not apply to our study, as we worked with species that maintain a constant level of attractiveness throughout the year

Citizen scientists have become an invaluable source of data for ecology and conservation efforts. However, to harness the full potential of these data, it is crucial to acknowledge their inherent limitations, including temporal biases in recording activity. In our study, we found that levels of recording activity by citizen scientists are higher on weekends, in days with milder temperatures, with little or no precipitation and wind, and predominantly in the spring. To effectively use citizen science data for ecological research, it is crucial to account for these habits. Robust statistical methods must be adopted to correct for these influences, ensuring a more accurate representation of ecological dynamics. For instance, adjusting for the increased recording activity during favourable weather conditions and specific times of the year can help mitigate the skewed data and provide a clearer picture of true ecological and biodiversity patterns.

## Author contributions

CC conceived the idea with input from all other authors, performed the analyses and contributed to the writing of the manuscript. PT collect the data and contribute to the writing of the manuscript. ITR and SC contributed to the writing of the manuscript.

## Availability of data and materials

Data and R code will be made publicly available upon publication of the manuscript.

